# Abnormal global longitudinal strain and reduced serum inflammatory markers in cardiac AL amyloidosis patients without significant amyloid fibril deposition

**DOI:** 10.1101/2024.03.14.584987

**Authors:** Camille V. Edwards, Grace M. Ferri, Josue Villegas-Galaviz, Sabrina Ghosh, Pushpinder Singh Bawa, Feiya Wang, Elena Klimtchuk, Tinuola B. Ajayi, Gareth J. Morgan, Tatiana Prokaeva, Andrew Staron, Frederick L. Ruberg, Vaishali Sanchorawala, Richard M. Giadone, George J. Murphy

## Abstract

**Background:** Cardiac dysfunction in AL amyloidosis is thought to be partly related to the direct impact of AL LCs on cardiomyocyte function, with the degree of dysfunction at diagnosis as a major determinant of clinical outcomes. Nonetheless, mechanisms underlying LC-induced myocardial toxicity are not well understood.

**Methods:** We identified gene expression changes correlating with human cardiac cells exposed to a cardiomyopathy-associated κAL LC. We then sought to confirm these findings in a clinical dataset by focusing on clinical parameters associated with the pathways dysregulated at the gene expression level.

**Results:** Upon exposure to a cardiomyopathy-associated κAL LC, cardiac cells exhibited gene expression changes related to myocardial contractile function and inflammation, leading us to hypothesize that there could be clinically detectable changes in GLS on echocardiogram and serum inflammatory markers in patients. Thus, we identified 29 patients with normal IVSd but abnormal cardiac biomarkers suggestive of LC-induced cardiac dysfunction. These patients display early cardiac biomarker staging, abnormal GLS, and significantly reduced serum inflammatory markers compared to patients with clinically evident amyloid fibril deposition.

**Conclusion:** Collectively, our findings highlight early molecular and functional signatures of cardiac AL amyloidosis, with potential impact for developing improved patient biomarkers and novel therapeutics.

## Introduction

Systemic immunoglobulin light chain (AL) amyloidosis is a disorder marked by the production of amyloidogenic immunoglobulin light chains (AL LCs) by clonal plasma cells within the bone marrow. [1] Disease pathology is manifested in several ways.

First, AL LCs form oligomers that can directly cause cardiac and renal dysfunction via proteotoxic effects. [1] LC oligomers also become amyloid fibrils that deposit in downstream target organs, with associated architectural distortion, structural damage, organ dysfunction, and eventually, organ failure. [1] Currently, the treatment of AL amyloidosis involves eradicating clonal plasma cells via chemoimmunotherapy, targeted therapies, or high-dose melphalan and autologous stem cell transplantation.. [1] Eradicating the plasma cell clone decreases the production of LCs which ceases further amyloidogenesis and, in some cases, permits a gradual but unpredictable regression of amyloid deposits. [2] While plasma cell-directed therapy improves organ function in some patients over time, the degree of cardiac dysfunction at diagnosis is a major predictor of morbidity, tolerance of therapies, and early mortality in patients with AL amyloidosis. [3–8] Heart failure results from a multi-step process involving cellular dysfunction directly induced by AL LCs with an LC sequence derived from particular germline gene donors (amyloidogenic, cardiomyopathy-associated LC) as well as disruption of normal myocardial structure from extracellular amyloid fibril deposition. [9–19] Evidence suggests that exposure to amyloidogenic, cardiomyopathy-associated LCs may represent the initial event leading to cardiac dysfunction in AL amyloidosis. [9,10,17] Additionally, the unique phenomenon of LC-induced cardiac dysfunction may contribute to the disproportionately inferior prognosis of cardiac AL amyloidosis over other cardiac amyloidoses (such as transthyretin-related amyloid cardiomyopathy) despite similar levels of amyloid infiltration. [19–21] While anti-fibril therapies are currently in development [22–25], no therapies are available to prevent LC-induced cardiac dysfunction. Moreover, the earliest events associated with cardiac dysfunction in AL amyloidosis remain elusive, hindering the development of novel biomarkers and therapies. Our group and others have shown that when cardiomyocytes are exposed to amyloidogenic, cardiomyopathy-associated LCs, high levels of reactive oxygen species are generated leading to mitochondrial damage, lysosomal dysfunction, and apoptosis, possibly through activation of the p38 mitogen-activated protein kinase (MAPK) pathway. [9–12]

Transcriptomic profiling can help further characterize pathological insults and improve biomarker discovery by allowing for the rapid and simultaneous measurement of thousands of variables associated with disease states. [26,27] However, applying laboratory-based indicators of cardiac dysfunction for clinical use is more complex.

Over the years, the availability of cardiac biomarkers (brain natriuretic peptide [BNP], N terminal prohormone of BNP [NT-proBNP] and Troponin I) has allowed us to stratify patients by clinical outcomes and has refined the approach to the management of AL amyloidosis. [28–30] Routine serial measurements of cardiac biomarkers are used to identify patients with AL amyloidosis who have developed overt organ damage and represent the standard of care for risk stratification for newly diagnosed AL amyloidosis. [28–31] However, cardiac biomarkers can be impacted by sodium balance, fluid shifts, diuretic use, and renal impairment. [32–36] As there is no standard method for confirming the presence of LC-induced cardiac dysfunction at diagnosis, the identification of unique clinical features associated with direct LC-induced cardiac dysfunction in patients without clinical evidence of significant amyloid fibril deposition is paramount as a next step in translating scientific discoveries to clinical practice.

Previously, we characterized neuronal SH-SY5Y and cardiac AC16 cells to an amyloidogenic, cardiomyopathy-associated kappa 1 LC (cardiomyopathy-associated κAL LC) as compared to amyloidogenic transthyretin (TTR) via RNA sequencing (RNAseq). [37] In doing so, we demonstrated that AC16 cells exposed to cardiomyopathy-associated κAL LC displayed a significantly higher number of differentially expressed genes than AC16 cells exposed to non-AL amyloidogenic proteins with no significant overlap in differentially regulated genes between study groups. [37] Herein, we used human induced pluripotent stem cell (iPSC)-derived cardiomyocytes to validate the gene expression changes of clinically relevant genes in cardiac dysfunction-related pathways that are differentially activated upon exposure to a cardiomyopathy-associated κAL LC. We then used this data to guide subsequent interrogation of our clinical dataset for measurable parameters that reflect gene expression changes observed in these cardiac dysfunction-related pathways. We focused our clinical analysis on a subset of patients with AL amyloidosis that demonstrate cardiac dysfunction without clinically apparent cardiac amyloid fibril deposition which we postulate to be the result of the direct myocardial toxic effects of amyloidogenic, cardiomyopathy-associated LCs.

## Materials and methods

### Protein preparation and characterization

Patient-derived kappa 1 immunoglobulin LC recombinant protein was prepared and generated as previously described. [38–40] Briefly, a bone marrow aspirate was obtained from a patient with an underlying kappa clonal plasma cell dyscrasia and associated cardiac involvement of AL amyloidosis. Red blood cells were lysed by treating the bone marrow aspirate with ammonium chloride. Thereafter, total RNA was extracted and used to synthesize complementary DNA which was then amplified by multiplex polymerase chain reaction, cloned, and sequenced as previously described. [38, 40] The LC was found to be derived from the IGKV1-33*01-IGKJ4*01-IGKC*01 germline gene donor. Nucleotide sequence was submitted to the GenBank under the accession number EF589383 with unique identifier AL-009L. Recombinant LC protein was generated via the transformation of DNA into E. coli using the heat shock method. In prior work, the resulting protein was purified, characterized by mass spectrometric analysis, and examined in various cardiac cell cultures. [38]

### Induced pluripotent stem cell (iPSC) generation and maintenance

As previously described, a wild-type non-diseased iPSC line BU5 was generated through hSTEMCCA lentiviral transduction of human peripheral blood mononuclear cells, meeting quality control criteria for pluripotency and functionality. [41, 42] iPSCs were maintained in mTESR (StemCell Technologies) on Matrigel (Corning Matrigel hESC-qualified Matrix; #354277) and passaged using ReLeSR (StemCell Technologies).

### Directed differentiation of iPSCs into cardiomyocytes

iPSCs were plated at 2 x 10^6^ viable cells per well on Matrigel-coated 6-well plates with 2mL of mTESR and incubated at 37 °C under normoxic, 5% carbon dioxide conditions. mTESR was replaced every 24 hours for two days. After two days, cardiomyocyte differentiation was induced using STEMdiff^TM^ Cardiomyocyte differentiation media [StemCell Technologies; #05010] only if > 95% confluence of iPSCs was achieved (Day 0). Day 8, 11, and 15 iPSC-derived cardiomyocytes were used for characterization assays by RT-qPCR and directly visualized via live cell imaging on the Keyence BZ- X710 All-in-one Fluorescence Microscope. Day 15 iPSC-derived cardiomyocytes, which demonstrated spontaneous beating, were used for dosing experiments.

### Dosing iPSC-derived cardiomyocytes with amyloidogenic, cardiomyopathy-associated kappa 1 LC

Spontaneously beating day 15 iPSC-derived cardiomyocytes were dosed with 0.02 mg/mL of recombinant kappa 1 LC derived from a patient with cardiac amyloidosis (cardiomyopathy-associated κAL LC). Undosed iPSC-derived cardiomyocytes, hereafter referred to as naïve cells, were used as a negative control. Dosed and naïve cells were incubated in triplicate at 37°C (normoxic and 5% carbon dioxide conditions) and observed intermittently over 48 hours via live cell imaging on the Keyence BZ- X710 All-in-one Fluorescence Microscope. Videos of 10 seconds were recorded at four time points (pre-dosing/ baseline, 6 hours, 24 hours, and 48 hours) for each sample, and beats were counted at baseline and 24 hours via direct visualization while cells were maintained at 37°C (normoxic and 5% carbon dioxide conditions). iPSC-derived cardiomyocytes were then harvested for flow cytometry or RNA extraction.

### RNA isolation, cDNA synthesis, and qRT-PCR

RNA was extracted from iPSC-derived cardiomyocytes and purified using a genomic DNA removal kit (ThermoFisher, Waltham, MA, Cat. no. AM1906). One microgram of purified RNA was used to generate cDNA using the High-Capacity cDNA Reverse Transcription Kit (Applied Biosystems, Foster City, CA, Cat. no. 43–688-13). The cDNA was PCR-amplified to add the sequencing adaptors. TaqMan^TM^ Universal Master Mix II, with UNG (ThermoFisher, Waltham, MA, Cat. no. 4440038), was used to perform qRT-PCR of cDNA. Samples were run in triplicate. TaqMan^TM^ Gene Expression Assays used include the following: *18S* [Hs99999901_s1], *MMP3* [Hs00968305_m1], *ROCK1* [Hs01127701_m1], *GPR183* [Hs00270639_s1], *CCL11* [Hs00237013_m1], *KDR* [Hs00911700_m1], *NKX2.5* [Hs00231763_m1], *TNNT2* [Hs00165960_m1]. The DDCT method was used to calculate fold change. Undetermined CT values were taken to be 40.

### Statistics

#### Transcriptomic data

Gene expression and pathway analyses were performed using data from Ghosh et al. 2022. [37] Functional predictions were made using the Enrichr gene set enrichment analysis web server [43–45] cross-referenced with the Gene Ontology Biological Process database [http://geneontology.org/]. The current version of the Broad Institute Connectivity Map [CMap] [46] was used to query the gene expression signature derived from bulk RNA sequencing on AC16 ventricular cardiomyocytes exposed to the cardiomyopathy-associated κAL LC. The Broad Institute CMap is a detailed catalog of transcriptional signatures derived from various cell types exposed to genetic or pharmacologic perturbagens. [46] An enrichment score was calculated for each condition related to each perturbagen-induced profile. The enrichment scores were then used to determine a connectivity score between -1 and 1 for each perturbagen-disease combination. The biologic activity of a given perturbagen [pharmacologic compound in this case] was determined by transcriptional activity score [TAS] and signature strength, where pharmacologic compounds with TAS > 0.2 and signature strength > 150 were considered biologically active.

#### Clinical data

Demographic, clinical, laboratory, and histological data were collected from either a prospectively maintained database at the Boston University Amyloidosis Center or the electronic medical record. In addition, patient charts were reviewed to ensure data completeness and accuracy. Echocardiographic variables at AL diagnosis, including left ventricular ejection fraction (LVEF) and global longitudinal strain (GLS), were collected from the electronic health record or post-processed manually using TomTec 2D cardiac performance analysis. Serum inflammatory markers at AL diagnosis, including C-reactive protein (CRP); erythrocyte sedimentation rate (ESR); lactate dehydrogenase (LDH); D-dimer, uric acid, and ferritin, were also collected from the electronic health record. The database was searched for patients with biopsy-proven AL amyloidosis consecutively evaluated in the Amyloid Clinic between January 1^st^, 2011 and March 31^st^, 2022. Patients with clinically suspected cardiac involvement (*n* = 426) were identified by the presence of elevated cardiac biomarkers per institutional reference ranges, including BNP > 72 mg/dL or NT-proBNP > 899 pg/mL or cardiac troponin I > 0.033 ng/mL. For the final analysis, we defined presumed LC-induced cardiac dysfunction as (1) elevated cardiac biomarkers, (2) normal echocardiographic intraventricular septal thickness (≤ 11 mm in males and ≤ 10 mm in females), (3) estimated glomerular filtration rate ≥ 60 mL/min/1.73m^2^, (4) absence of late gadolinium enhancement on cardiac magnetic resonance imaging (if performed), and (5) negative endomyocardial biopsy (if performed). Patients were excluded from the analysis if they met the following criteria (1) increased echocardiographic intraventricular septal thickness (> 11 mm in males and > 10 mm in females), (2) abnormal cardiac magnetic resonance imaging (MRI) suggestive of cardiac amyloidosis, (3) endomyocardial biopsy-proven cardiac amyloid infiltration, or (4) estimated glomerular filtration rate (eGFR) < 60 mL/min/1.73m^2^ that could increase the concentration of cardiac biomarkers. Descriptive statistics, including mean, median, and interquartile ranges, were used to summarize baseline demographic and clinical characteristics.

### Study approval

All patients provided written informed consent for using their medical records for research per the Declaration of Helsinki. The study was approved by the Institutional Review Board at Boston University [**ClinicalTrials.gov identifier:** <u>NCT00898235</u>].

### Data availability

The data that support the findings of this study are available on request, as applicable, from the corresponding author, CVE. The clinical data are not publicly available due to containing information that could compromise the privacy of research participants.

## Results

### AC16 cells exposed to an amyloidogenic, cardiomyopathy-associated kappa 1 light chain exhibit differential expression of genes associated with cardiac dysfunction

Our group previously examined transcriptomic changes occurring in AC16 human ventricular cardiomyocyte cells exposed to a cardiomyopathy-associated κAL LC as described above from the Boston University Amyloidosis Center and amyloidogenic transthyretin (TTR) proteins [37]. Here, we performed a post-hoc functional gene set enrichment analysis (fgsea) to evaluate hallmark pathways enriched upon exposure of AC16 cells to a cardiomyopathy-associated κAL LC. This analysis revealed the unique enrichment of pathways related to (1) cardiac muscle hypertrophy, (2) cardiac muscle contraction and relaxation, and (3) regulation of heart rate by cardiac conduction. We also noted the downregulation of hallmark pathways related to (1) inflammatory responses, (2) adaptive immune responses, and (3) oxidative stress-induced cell death in AC16 cells exposed to a cardiomyopathy-associated κAL LC compared to cardiomyopathy-associated TTR proteins (**Figure 1A**). For validation of previously reported transcriptomic changes, we selected significantly differentially expressed genes that are relevant to the pathophysiology of cardiomyopathies, and related to the hallmark pathways uniquely dysregulated in our dataset (*MMP3, ROCK1, GPR183,* and *CCL11*) (**Supplemental Figure 1**). [47–60]

**Figure 1.**
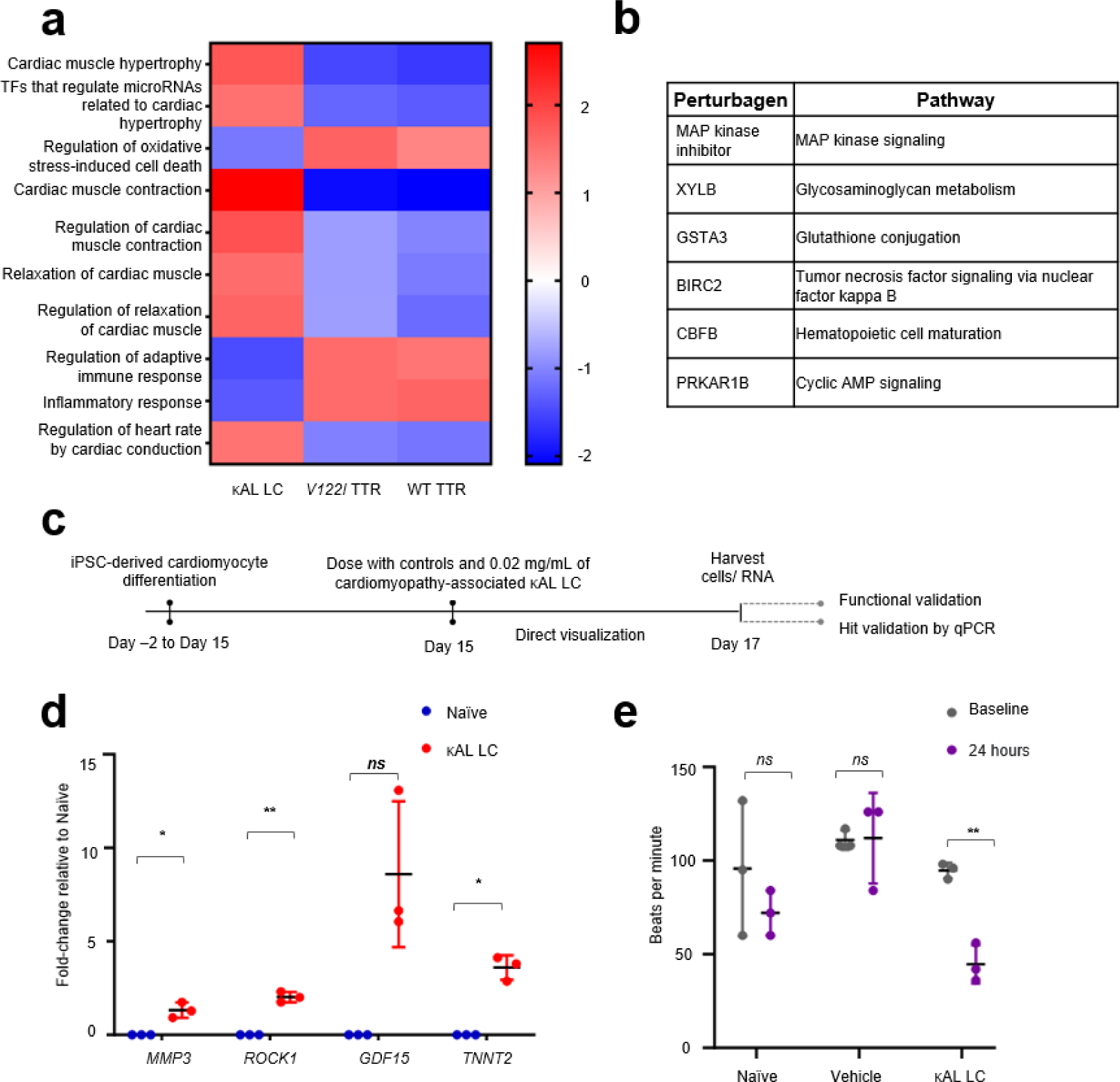
Transcriptional and functional response of human cardiac cells exposed to an amyloidogenic, cardiomyopathy-associated kappa 1 LC. (a) Post-hoc analysis of RNAseq data from Ghosh et al.^36^ using functional gene set enrichment analysis to compare the top hallmark pathways dysregulated in AC16 cells exposed to a cardiomyopathy-associated κAL LC versus TTR proteins. (b) Top six biologically active pharmacologic compounds that reverse the transcriptional signature of LC-induced cardiac dysfunction as determined by a connectivity map (CMap) analysis of differentially expressed genes in AC16 cells exposed to a cardiomyopathy-associated κAL LC protein, *p* < 0.05. The biologic activity was determined by a high transcriptional activity score > 0.2 and high signature strength > 150. (c) Summary of experimental design for dosing iPSC-derived cardiomyocytes with a cardiomyopathy-associated κAL LC (0.02 mg/mL) derived from a patient with cardiac AL amyloidosis. Cells were intermittently observed via live cell imaging for 48 hours. RNA was then isolated from each condition for qRT-PCR validation of select target genes. (d) qRT-PCR validation of select target genes related to hallmark pathways (*MMP3*, *ROCK1, GDF15*, *TNNT2*) in iPSC-derived cardiomyocytes exposed to a cardiomyopathy-associated κAL LC protein. (e) Beating of iPSC-derived cardiomyocytes exposed to a cardiomyopathy-associated κAL LC protein. Dot plots display error bars. Each *n* represents a set of assays with at least 3 technical replicates from a distinct biological replicate. * p < 0.05, ** p < 0.01, *** p < 0.001, ns - not significant; unpaired t-tests. Abbreviations: TFs: transcription factors; κAL LC: amyloidogenic, cardiomyopathy-associated kappa 1 LC protein; TTR: transthyretin amyloid; WT TTR: wild-type TTR protein; V122I: genetic variant of cardiomyopathy-associated TTR protein; TAS: transcriptional activity score; XYLB: xylulokinase; GSTA3: Glutathione S-transferase alpha 3; BIRC2: baculoviral IAP repeat containing 2; CBFB: core-binding factor subunit beta; PRKAR1B: protein kinase cAMP-dependent type I regulatory subunit beta. Naïve: cardiomyocytes in respective culture media; Vehicle: 70% ethanol.

Significantly differentially expressed genes (FDR <0.05) were then analyzed using the connectivity map (CMap) to identify pharmacologic compounds that may resolve the transcriptional signatures noted with cardiomyopathy-associated κAL LC exposure. To this end, we identified six compounds targeting pathways related to LC-mediated cardiac dysfunction, including: (1) mitogen-activated protein kinase (MAPK), (2) glycosaminoglycan metabolism, (3) glutathione conjugation, (4) tumor necrosis factor signaling via nuclear factor kappa B, (5) hematopoietic cell maturation, and (6) cyclic adenosine monophosphate (cAMP) signaling pathways (**Figure 1B**)

### iPSC-derived cardiac cells exposed to an amyloidogenic, cardiomyopathy-associated kappa 1 light chain exhibit transcriptional and functional changes suggestive of cardiac dysfunction

We next sought to confirm the differential expression of genes identified in our previously published RNAseq dataset in more physiologically relevant cardiac cells using human iPSC-derived cardiomyocytes exposed to the same cardiomyopathy-associated κAL LC (**Figure 1C**, **Supplemental Figure 2**). For validation of previously reported transcriptomic changes, we selected significantly differentially expressed genes related to clinically relevant phenomena such as adaptive cardiac remodelling (*MMP3, ROCK1, GDF15*) [47–54, 61] and with functional relevance to oxidative stress-induced cardiomyocyte damage (*GDF15*, *TNNT2*) [61–64]. Expression of protein-coding genes related to adaptive cardiac remodelling, and oxidative stress-induced cardiomyocyte damage were significantly upregulated in iPSC-derived cardiomyocytes exposed to a cardiomyopathy-associated κAL LC (**Figure 1D**).

Since pathways related to the regulation of heart rate were enriched in AC16 cells exposed to a cardiomyopathy-associated κAL LC (**Figure 1A**), changes in the rhythmic beating of iPSC-derived cardiomyocytes were also assessed (**Supplemental Video file 1**). Notably, the cardiomyopathy-associated κAL LC induced negative chronotropic effects on iPSC-derived cardiomyocytes within 24 hours of exposure, as evidenced by a statistically significant reduction in beats per minute (**Figure 1E**).

### Transcriptomic data guides clinical analyses in a novel clinical dataset highlighting features of light chain-induced cardiac dysfunction without clinical evidence of amyloid fibril deposition

Since genes related to regulating cardiac muscle contraction were dysregulated in the transcriptomic dataset, we hypothesized that GLS (an early echocardiographic measure of myocardial contractility) abnormalities would be detectable in an AL cohort with suspected LC-induced cardiac dysfunction. Thus, we retrospectively evaluated the baseline clinical and echocardiographic features of the 29 patients identified with what we propose to be LC-induced cardiac dysfunction. Baseline patient characteristics are summarized in **Table 1**. The majority of patients were male (55%; *n* = 16). The median age of patients was 65 years (range, 46-83). LC isotype was lambda in 69% (*n* = 20) and kappa in 31% (*n* = 9) of patients, respectively. The difference in free light chains (dFLC) was ≥ 50 mg/L in 62% (*n* = 18) of patients at diagnosis. Patients also had early cardiac disease with median BNP of 237 pg/mL and median troponin I of 0.048 ng/L. Notably, 76% (*n* = 22) of patients had Boston University Stage II cardiac disease. [65] GLS data were available in 16 cases. The median GLS value was -15.5% (IQR -7 to - 21.7) compared to the normal value of ≤ -18%. An abnormal GLS was observed in 13/16 (81%) patients (**Figures 2**). Notably, LVEF, a global measure of cardiac systolic function, was similar in patients with normal and abnormal GLS values.

**Figure 2:**
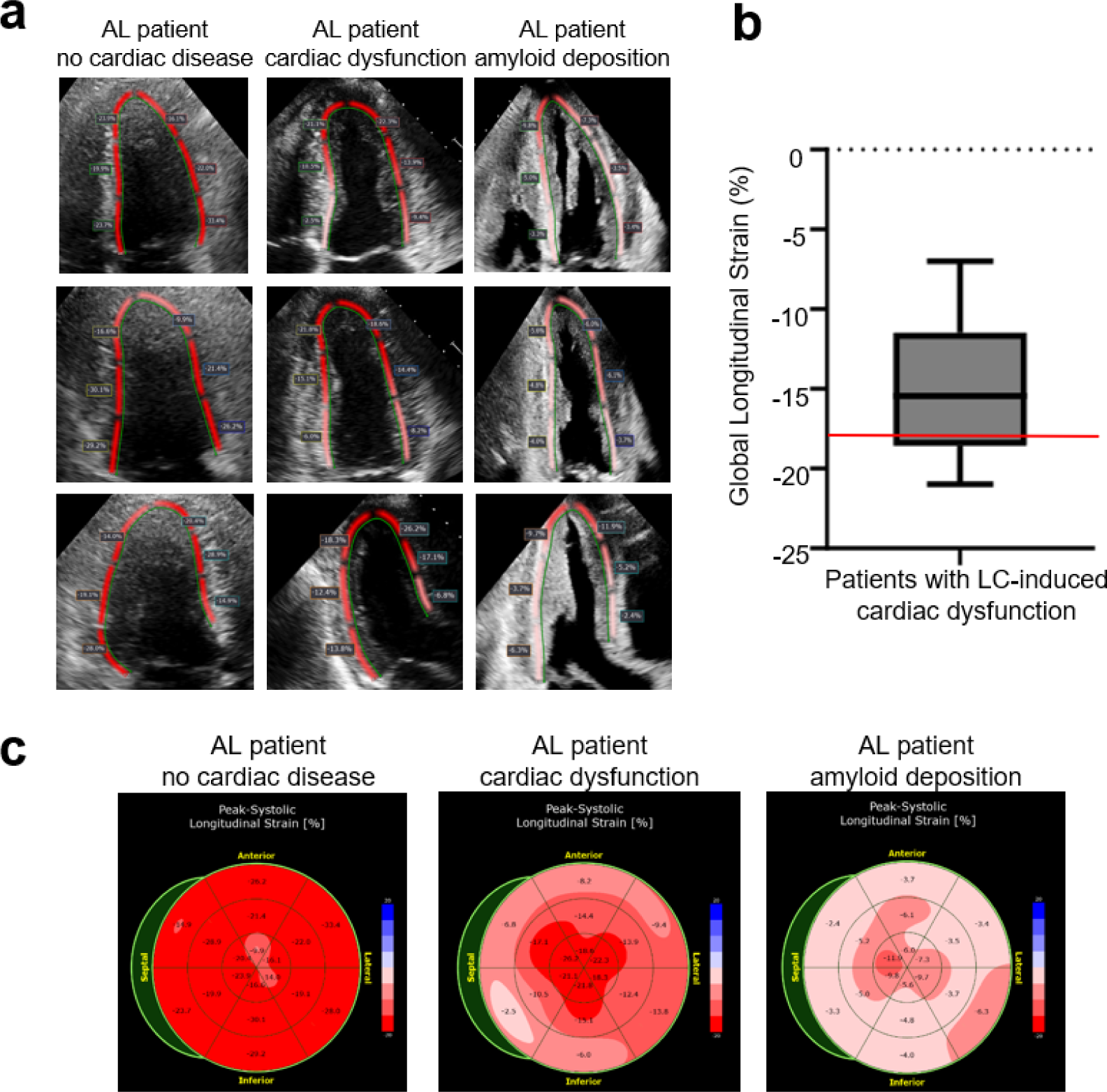
Cardiac AL impacts global longitudinal strain on echocardiography. (a) End-systolic speckle tracking images for three cardiac windows in patients with cardiac AL amyloidosis. Images from top to bottom are apical four-chamber view, apical two- chamber view, apical three-chamber view [1] no cardiac disease and normal GLS (- 21.4%), [2] LC-induced cardiac dysfunction and slightly depressed GLS (-14.3%), and [3] cardiac involvement and amyloid fibril deposition with severely depressed GLS (- 5.7%). **(b)** Global longitudinal strain among 16 patients with AL amyloidosis and LC- induced cardiac dysfunction. The median GLS value was -15.5% (IQR -7 to). Box plot displays median (-15.5%), 25^th^ quartile (-18.5%), and 75^th^ quartile (-11.9%), minimum value (-21.7%), and maximum value (-7%) for GLS. Normal is defined as ≤ -18.0% (red line), thus more negative is more normal and less negative (approaching 0, dotted line) is more abnormal. **(c)** Global longitudinal strain map in three patients with AL amyloidosis with [1] no cardiac disease and normal GLS (-21.4%), [2] LC-induced cardiac dysfunction and slightly depressed GLS (-14.3%), and [3] cardiac am significant fibril deposition with severely depressed GLS (-5.7%). **Abbreviations:** GLS: global longitudinal strain; LC: light chain

**Table 1:** Baseline Demographic, Clinical, and Echocardiographic Features of 29 Patients with AL amyloidosis and LC-induced Cardiac Dysfunction.

|  |  |
| --- | --- |
| **Demographics and clinical characteristics** | ***n* = 29** |
| Age at diagnosis years, median (IQR) | 65 (46 – 83) |
| Male sex, n (%) | 16 (55) |
| **Diagnosis** |  |
| AL, n (%) | 28 (96.6) |
| MMAL, n (%) | 1 (3.4) |
| **Light chain restriction type** |  |
| Lambda, n (%) | 20 (69) |
| Kappa, n (%) | 9 (31) |
| **dFLC** |  |
| 10 - < 50 mg/L, n (%) | 11 (38) |
| ≥50, n (%) | 18 (62) |
| **Cardiac biomarkers and echocardiographic features** |  |
|  | 237 (82 - 1932) |
| BNP pg/mL, median (IQR) | 0.048 (0.006 – 1.217) |
| Troponin I ng/L, median (IQR) | 62 (30 – 76) |
| LVEF (%), median (IQR) |  |
| **Boston University Cardiac Stage<sup>A</sup>** |  |
| Stage II, n (%) | 22 (76) |
| Stage IIIA, n (%) | 5 (17) |
| Stage IIIB, n (%) | 2 (7) |
| <sup>A</sup>Stage I not listed. Patients with normal cardiac biomarkers were excluded from this cohort. AL – AL amyloidosis. MMAL – multiple myeloma-associated AL amyloidosis. dFLC – difference in involved and uninvolved free light chain concentration. BNP – brain natriuretic peptide (normal range: 0 – 72.3 pg/mL). Troponin I normal range: 0.01 – 0.04 ng/L. LVEF – left ventricular ejection fraction (normal range 55 – 70%). IQR - interquartile range. |  |

Moreover, genes related to the regulation of inflammatory responses were downregulated in our transcriptomic dataset. As a result, we hypothesized that there could be clinically measurable differences in serum inflammatory markers in patients with presumed LC-induced cardiac dysfunction as compared to those with clinically evident amyloid fibril deposition. In accordance, when compared to patients with clinically evident amyloid fibril deposition on echocardiogram, patients with presumed LC-induced cardiac dysfunction had significantly lower mean serum inflammatory markers, including CRP (mean 10.22 [0.1-295.3] versus 6.007 [0.3-28.8]), D-dimer (mean 874 [152–11580] versus 517 [157–1304]), LDH (mean 287 [102–860] versus 262 [163–463]), and uric acid (mean 7.42 [2.30-18.00] versus 5.97 [2.80-9.30]) (**Figure 3**).

**Figure 3:**
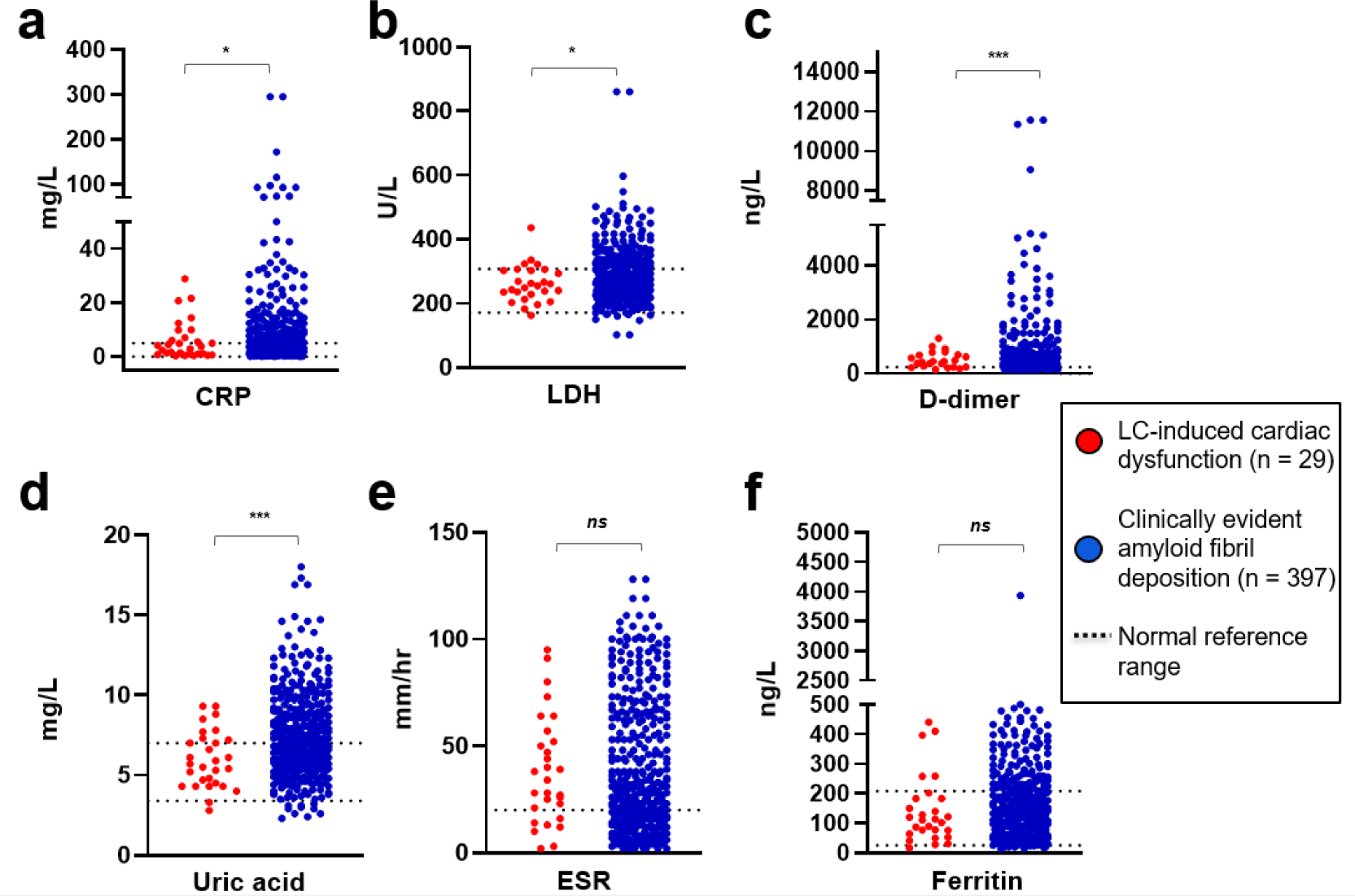
Patients with light chain-induced cardiac dysfunction exhibit lower serum inflammatory markers than those with clinically evident amyloid fibril deposition. We identified 426 cardiac AL patients and stratified the cohort based on the absence or presence of increased wall thickness on echocardiogram. Presumed LC-induced cardiac dysfunction (*n* = 29, red scatter plots) was defined as (1) elevated cardiac biomarkers (BNP > 72 pg/mL or NT-proBNP > 899 pg/mL or troponin I > 0.033 ng/mL, per institutional reference ranges), (2) normal echocardiographic intraventricular septal thickness (≤ 11 mm in males and ≤ 10 mm in females), (3) estimated glomerular filtration rate ≥ 60 mL/min/1.73m^2^, (4) absence of late gadolinium enhancement on cardiac magnetic resonance imaging (if performed), and (5) negative endomyocardial biopsy (if performed). Patients with significant clinical evidence of cardiac amyloid fibril deposition (*n* = 397, blue scatter plots) had significantly higher serum inflammatory markers (mean, range): (a) CRP (mean 10.22 [0.1-295.3] versus 6.007 [0.3-28.8]), (b) LDH (mean 287 [102–860] versus 262 [163–463]), (c) D-dimer (mean 874 [152–11580] versus 517 [157–1304]), and (d) uric acid (mean 7.42 [2.30-18.00] versus 5.97 [2.80-9.30]). The groups did not significantly differ in (e) ESR (mean 39.25 [2.00-128.00] versus 38.48 [2.00- 95.00]) and (f) ferritin (mean 272 [13–3933] versus 190 [17–1497]). * p < 0.05, ** p < 0.01, *** p < 0.001, ns - not significant; unpaired t-tests. Reference ranges (black dotted lines): CRP (C-reactive protein) 0 – 5 mg/L, erythrocyte sedimentation rate (ESR) 0 – 20 mm/hr, D-dimer < 243 ng/mL, ferritin 26 – 209 ng/mL, uric acid 3.4 – 7.0 mg/dL, lactate dehydrogenase (LDH) 171 – 308 U/L.

## Discussion

Here, we utilized a previously generated transcriptomic dataset to evaluate dysregulated pathways upon exposure of immortalized cardiac AC16 cells to a cardiomyopathy-associated κAL LC. We then employed more physiologically relevant, human iPSC-derived cardiomyocytes to confirm changes in the expression of clinically relevant genes. Finally, we sought to identify a clinical correlate for our transcriptomic findings by searching a clinical database for AL patients with evidence of LC-induced cardiac dysfunction and evaluating serum and echocardiographic parameters in this unique cohort. Altogether, our results suggest that on a transcriptomic level, cardiomyocytes display dysregulation of cardiac contractility upon exposure to a cardiomyopathy-associated κAL LC which could manifest clinically as early cardiac biomarker stage disease and abnormal GLS in a subset of AL patients. We also found that dysregulation of inflammatory responses in cardiomyocytes exposed to cardiomyopathy-associated LCs could potentially be reflected as clinically measurable differences in serum inflammatory markers in patients with presumed LC-induced cardiac dysfunction versus clinically evident cardiac amyloid fibril deposition. We postulate that these molecular and functional signatures directly result from the proteotoxic effects of an amyloidogenic, cardiomyopathy-associated LC (LC-induced cardiac dysfunction).

At the transcriptional level, cardiac cells exhibit the significant upregulation of hallmark pathways that control cardiac hypertrophy upon exposure to a cardiomyopathy-associated κAL LC. We postulate that this transcriptional pattern could provide evidence for an adaptive hypertrophic phenotype that would reduce oxygen consumption, diminish cardiomyocyte stress, and preserve overall cardiac function [66,67] in the initial phases of exposure to cardiomyopathy-associated κAL LC. Moreover, it is known that compensatory cardiac hypertrophy becomes maladaptive due to specific insults, including oxidative stress, inflammation, or mechanical stress, which further leads to cardiac remodeling, contractile dysfunction, and eventually heart failure. [47, 49–54, 61, 67, 68, 69] Cardiac remodeling is characterized by molecular, cellular, and structural changes that clinically manifest as changes in heart size, shape, and function. [70] In line with this knowledge, subsequent validation of our transcriptional signature by RT-qPCR confirmed the upregulation of genes involved in cardiac remodeling and oxidative stress-induced cell death (*MMP3*, *ROCK1*, *GDF15*, *TNNT2*).

Importantly, the genes validated (*MMP3*, *ROCK1*, *GDF15*, *TNNT2*) encode for proteins that are measurable in patient sera during myocardial injury and maladaptive cardiac remodeling. [47, 49–54, 61, 67, 68, 69] Our group has previously shown that increasing serum levels of metalloproteinases (MMPs [coded by *MMP* genes]) and tissue inhibitors of MMPs (TIMPs) correlate with echocardiographic features such as left ventricular wall thickness, left ventricular mass, and diastolic dysfunction in AL amyloidosis to a much greater degree than in non-AL cardiac amyloidosis. [53, 54] Additionally, GDF15 has been found to improve prognostication of cardiovascular diseases independently and in combination with traditional cardiac biomarkers (such as BNP, NT-proBNP, and troponin I). [61, 71–73] While measuring BNP, NT-proBNP, and troponin I is standard for risk stratification of patients with newly diagnosed AL amyloidosis, serum concentrations can be impacted by fluid shifts and renal impairment, suggesting a need for complementary biomarkers. Therefore, further exploration of these proteins (MMP3, ROCK1, GDF15) in AL patients with LC-induced cardiac dysfunction should be considered.

Additionally, our observation of dysregulated genes responsible for cardiac muscle contractility upon exposure to a cardiomyopathy-associated κAL LC prompted us to search clinical data for early echocardiographic changes in myocardial contractile function in cases with presumed LC-induced cardiac dysfunction. GLS is an early measure of myocardial contractility that is more sensitive and specific than the more widely used left ventricular ejection fraction. [74–76] In evaluating GLS, we found that the majority of patients with LC-induced cardiac dysfunction (81%, *n* = 13/16) displayed clinically detectable abnormalities in GLS. Abnormalities in GLS support conclusions proposed in previous studies that there are possibly subtle functional changes in myocardial contractility due to cardiomyopathy-associated AL LCs prior to overt systolic dysfunction. [9]

Interestingly, we noted the downregulation of inflammatory response genes in cardiomyocytes upon exposure to AL LCs. In contrast, Jordan et al. reported the upregulation of cytokines and chemokines in human cardiomyocytes exposed to amyloid fibrils, suggesting the initiation of an inflammatory response to amyloid fibrils. [77] Here, we noted clinically measurable differences in serum inflammatory markers that are potentially congruent with these findings. Specifically, in line with transcriptomic data, patients with LC-induced cardiac dysfunction had significantly lower serum inflammatory markers. It is unclear whether these responses are compensatory or unfavorable; however, these findings may indicate unique cellular responses to destabilized LCs versus amyloid fibrils.

Although our transcriptomic analysis reveals unique gene expression changes in cardiomyocytes exposed to a cardiomyopathy-associated κAL LC, it is difficult to ascertain the functional role of differentially expressed genes, particularly related to cardiac muscle cell contraction and negative regulation of heart rate. Additional phenotypic assays, including proteomics and electrophysiology and cardiac imaging, may shed light on this. Moreover, and importantly, given the genetic and structural diversity of AL LCs [38], the transcriptional signatures and clinical metrics described herein may not be identical for all cardiomyopathy-associated AL LCs. In this report, we focused on molecular changes in cardiac cells exposed to a single and well-characterized cardiomyopathy-associated κAL LC [38–40].

We observed that cardiomyocytes exhibit upregulation of cardiac muscle hypertrophy, cardiac muscle contractility, and oxidative stress-induced cardiomyocyte damage upon exposure to a cardiomyopathy-associated κAL LC protein. These findings could reflect an initiation of early cardiac dysfunction and maladaptive cardiac remodelling. Through combining transcriptomic and clinical data, we identified a previously undefined and uncharacterized subset of patients with AL amyloidosis who have: (1) early cardiac biomarker stage disease, (2) clinically detectable impairments in myocardial contractile function via GLS without significant clinical evidence of amyloid fibril deposition, and (3) clinically measurable differences in serum inflammatory markers, all of which we postulate could be the result of the direct myocardial effects of cardiomyopathy-associated AL LCs. As cardiac AL amyloidosis is often underdiagnosed due non-specific symptoms, the ability to detect early signatures of cardiac AL using readily available clinical parameters is paramount. In particular, with the advent of artificial intelligence and machine learning algorithms for diagnosing cardiac amyloidosis [78, 79] and light chain-mediated toxicity in AL [80], our studies support further exploration of GLS as an imaging biomarker combined with serum biomarkers for early diagnosis of cardiac AL.

## Acknowledgments

The Amyloidosis Center clinical database and samples repository are supported by the Amyloid Research Fund of Boston University Chobanian & Avedisian School of Medicine.

## Author contributions

C.V.E., R.M.G., and G.J. Murphy designed the study, devised experiments, analyzed data, and wrote the manuscript. G.J. Morgan and E.K. contributed to the study design and supplied valuable reagents. T.P. provided kappa 1 LC sequence information and contributed to the study design. V.S. and F.L.R. contributed to the study design. C.V.E. performed the experiments. G.M.F. collected, reviewed and analyzed clinical data. S.G., P.S.B., and F.W. analyzed the laboratory data. J.V.G., T.B.A., A.S., and F.L.R. analyzed clinical data. All authors provided feedback and assisted in writing the manuscript.

---

**Supplemental Figure 1.**
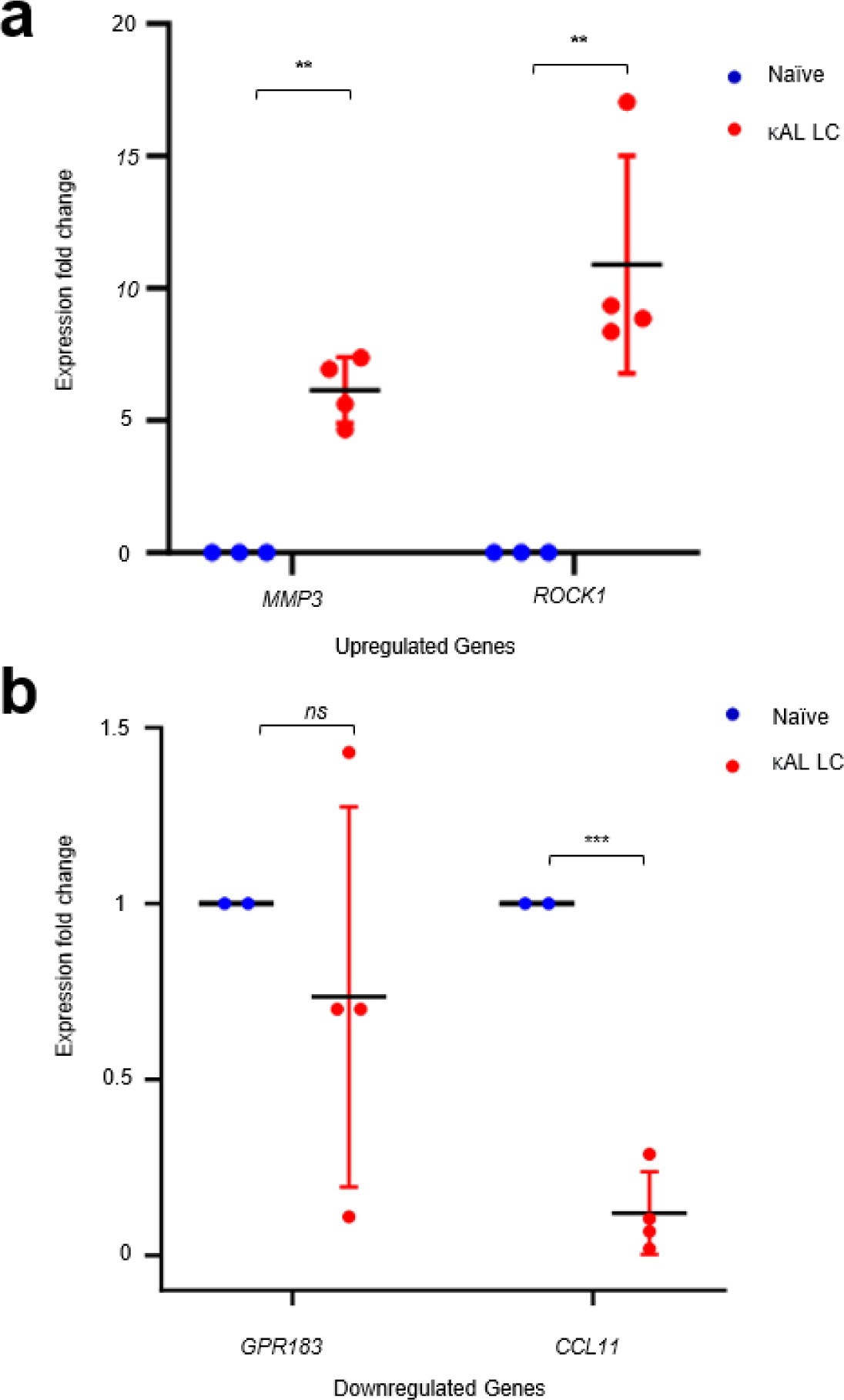
Transcriptional validation of genes dysregulated in AC16 human ventricular cardiomyocytes exposed to cardiomyopathy-associated κAL LC. qRT-PCR validating select target genes related to **(a)** upregulated hallmark pathways, cardiac remodeling (*MMP3*, *ROCK1*), and **(b)** downregulated hallmark pathways – inflammatory (*GPR183*) and adaptive immune responses (*CCL11*) in AC16 human ventricular cardiomyocytes exposed to cardiomyopathy-associated κAL LC. Each *n* represents a set of assays with at least 3 technical replicates from a distinct biological replicate. * *p < 0.05*, ** *p* < 0.01, *** *p* < 0.001, *ns - not significant* using unpaired t-test.

**Supplemental Figure 2.**
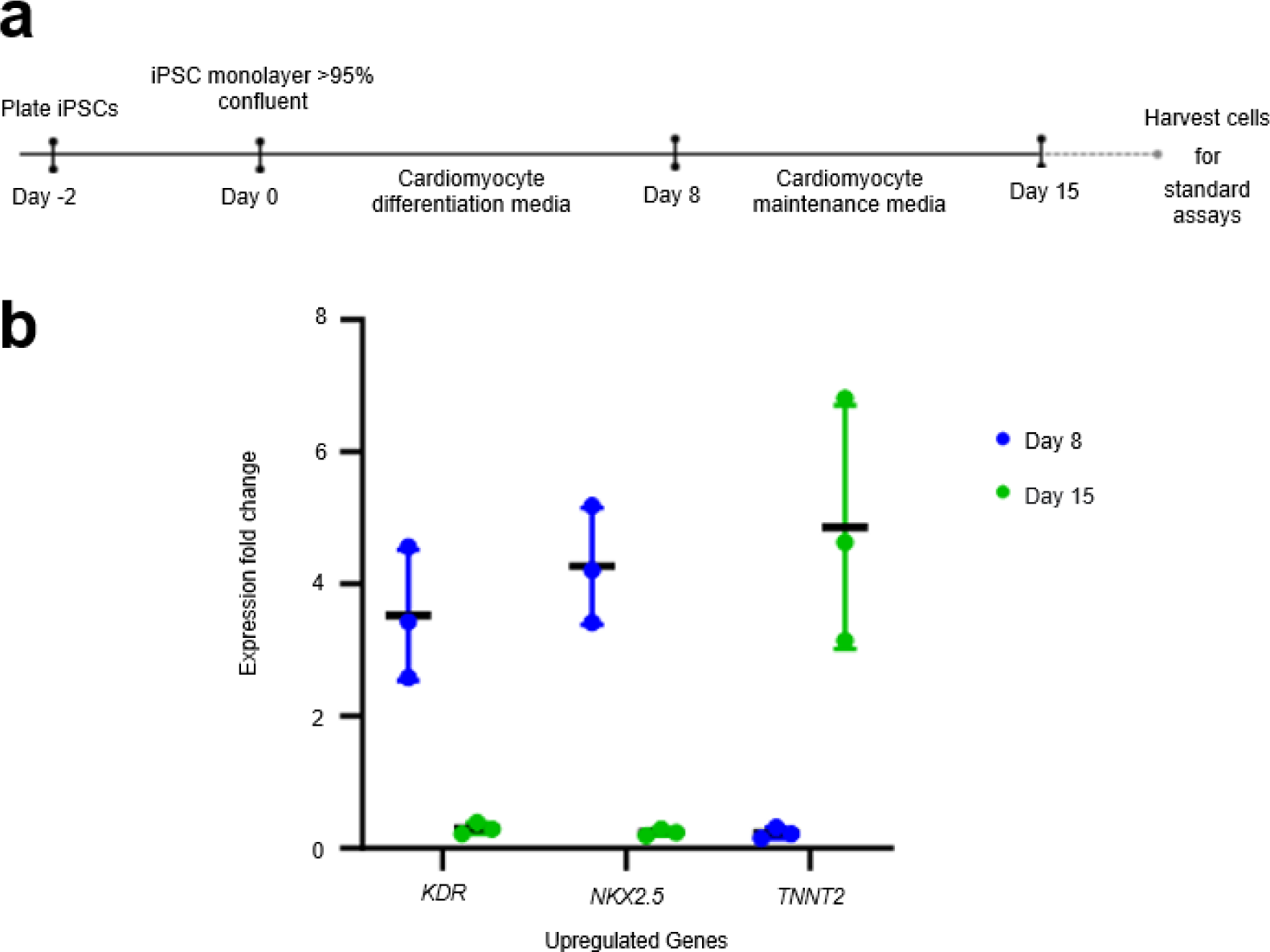
Characterization of iPSC-derived cardiomyocytes by q- RT-PCR and flow cytometry. **(a)** Cardiomyocyte differentiation was induced using STEMdiff^TM^ Cardiomyocyte differentiation media and cells were utilized 15 days later. **(b)** qRT-PCR to validate select markers for iPSCs (*NKX2.5*), cardiac mesoderm (*KDR*), and iPSC-derived cardiac cells (*TNNT2*) at Day 8 and Day 15 of the differentiation protocol. Each *n* represents a set of assays with at least 3 technical replicates from a distinct biological replicate. * *p < 0.05*, ** *p* < 0.01, *** *p* < 0.001, *ns - not significant* using unpaired t-test.

